# Recent zoonotic spillover and tropism shift of a Canine Coronavirus is associated with relaxed selection and putative loss of function in NTD subdomain of spike protein

**DOI:** 10.1101/2021.11.15.468709

**Authors:** Jordan D. Zehr, Sergei L. Kosakovsky Pond, Darren P. Martin, Kristina Ceres, Gary R. Whittaker, Laura B. Goodman, Michael J. Stanhope

**Affiliations:** Institute for Genomics and Evolutionary Medicine, Temple University, Philadelphia, PA, 19122, USA; Computational Biology Division, Department of Integrative Biomedical Sciences, Institute of Infectious Diseases and Molecular Medicine, University of Cape Town, Observatory, Cape Town, South Africa; Department of Population Medicine and Diagnostic Sciences, Cornell University, Ithaca, NY, 14853, USA; Department of Microbiology and Immunology, Cornell University, Ithaca, NY, 14853, USA; Baker Institute for Animal Health, Cornell University, Ithaca, NY, 14850, USA

## Abstract

A recent study reported the occurrence of Canine Coronavirus (CCoV) in nasopharyngeal swabs from a small number of patients hospitalized with pneumonia during a 2017-18 period in Sarawak, Malaysia. Because the genome sequence for one of these isolates is available, we conducted comparative evolutionary analyses of the spike gene of this strain (CCoV-HuPn-2018), with other available *Alphacoronavirus* 1 spike sequences. The most N-terminus subdomain (0-domain) of the CCoV-HuPn-2018 spike protein has sequence similarity to Transmissible Gastroenteritis Virus (TGEV) and CCoV2b strains, but not to other members of the type II Alphacoronaviruses (i.e., CCoV2a and Feline CoV2-FCoV2). This 0-domain in CCoV-HuPn-2018 has evidence for relaxed selection pressure, an increased rate of molecular evolution, and a number of unique amino acid substitutions relative to CCoV2b and TGEV sequences. A region of the 0-domain determined to be key to sialic acid binding and pathogenesis in TGEV had clear differences in amino acid sequences in CCoV-HuPn-2018 relative to both CCoV2b (enteric) and TGEV (enteric and respiratory). The 0-domain of CCoV-HuPn-2018 also had several sites inferred to be under positive diversifying selection, including sites within the signal peptide. Downstream of the 0-domain, FCoV2 shared sequence similarity to the CCoV2b and TGEV sequences, with analyses of this larger alignment identifying positively selected sites in the putative Receptor Binding Domain (RBD) and Connector Domain (CD). Recombination analyses strongly implicated a particular FCoV2 strain in the recombinant history of CCoV-HuPn-2018 with molecular divergence times estimated at around 60 years ago. We hypothesize that CCoV-HuPn-2018 had an enteric origin, but that it has lost that particular tropism, because of mutations in the sialic acid binding region of the spike 0-domain. As selection pressure on this region was reduced, the virus evolved a respiratory tropism, analogous to other *Alphacoronavirus* 1, such as Porcine Respiratory Coronavirus (PRCV), that have lost this region entirely. We also suggest that signals of positive selection in the signal peptide as well as other changes in the 0-domain of CCoV-HuPn-2018 could represent an adaptive role in this new host and that this could be in part due to the different spatial distribution of the N-linked glycan repertoire for this strain.

## 1. Introduction

The ongoing coronavirus (CoV) disease 19 (COVID-19) is the third documented animal to human CoV spillover, (SARS-CoV; SARS-CoV-2 and MERS-CoV), within the past two decades, to have resulted in a major epidemic. Coronaviruses (CoVs) that infect mammals (with the exception of pigs) belong principally to two genetic and serologic groups: the *Alphacoronavirus* (α) and *Betacoronavirus* (β) genera. *Alphacoronavirus* 1 is a species within the *Alpha* genus which comprises viruses that infect dogs, cats and pigs, and is further subdivided into type I and II based on serological parameters and genetic differences in the spike gene, although further genetic and biological differences are also apparent (see for e.g., CCoV1; Decaro et al. 2015). Vlasova et al. (2021) recently reported on an *Alphacoronavirus* 1 CoV resembling Canine CoV (CCoV), isolated from nasopharyngeal swabs of a small number of patients (8/301; seven of these eight were children), in Sarawak Malaysia, hospitalized with pneumonia during a 2017-18 period. The genome sequence of one of these isolates was obtained, while the other seven were diagnosed based on PCR tests. Genomically, the virus resembles a CCoV type II, but also shares high nucleotide sequence similarity with other type II *Alphacoronavirus* 1 viruses: feline CoV (FCOV2), and the porcine Transmissible Gastroenteritis Virus (TGEV).

Zoonotic transmissions of CoVs represent an important threat to human health, with many unknown possible reservoir hosts. The Vlasova et al. report is the first example of a Canine CoV isolated from a human patient with pneumonia, although the virus has not been confirmed as the causative pathogen and inter-individual transmission has not been established. However, the obvious importance of a jump from a companion animal such as a dog or a cat, or from a farm animal such as a pig, to a human host, raises concerns and questions about possible adaptative tropism of veterinary coronaviruses to humans. A key component of CoV host tropism is the binding of the spike protein with host cellular receptors. The spike protein is responsible for host receptor binding and fusion of the virus and host cell membranes (Li, 2016). It is comprised of the N-terminal S1 region, containing the receptor binding domain (RBD), and the C-terminal S2 region, responsible for membrane fusion. The CCoV receptor in dogs, and for the rest of the type II members of the *Alphacoronavirus* 1 species, is amino peptidase N (APN) (reviewed in Millet et al. 2021). The human common-cold coronavirus, HCoV-229E, another *Alphacoronavirus*, (not *Alphacoronavirus* 1) also uses APN. Structural studies involving porcine respiratory coronavirus (PRCV; closely related to TGEV) indicate that it binds to a site on porcine APN that differs from the site at which HCoV-229E binds to hAPN (Reguera et al. 2012; Wong et al. 2017), implying that there are multiple ways for this interaction to take place. Importantly, feline APN can serve as a functional receptor of type II CCoV, TGEV and human coronavirus HCoV-229E (Tresnan, Levis and Holmes 1996). While definitive experimental data are still lacking, the possibility of co-infections in cats implies that individual cells can become infected with these different CoVs, which could, in turn, generate novel recombinant strains. Spike gene recombination has played an important role in the evolution of the *Alphacoronavirus* 1 type II CoVs, involving recombination between dog, cat and pig viruses, including the complete replacement of the most N-terminus subdomain of CCoV2 with that of TGEV - an important event in the formation of CCoV2b (Wesley 1999). In keeping with this aspect of the *Alphacoronavirus* 1 group’s history, Vlasova et al., provide evidence that CCoV-CCoV-HuPn-2018 carries a recombinant CCoV/FCoV spike gene.

In addition to APN, alternative attachment factors and/or co-receptors are also known for some *Alphacoronavirus* 1 CoVs, including C-type lectins dendritic cell-specific intercellular adhesion molecule-3-grabbing non-integrin (DC-SIGN), heparan sulfate (HS), and sialic acid (reviewed in Millet et al. 2021). Thus, there are numerous possible avenues for developing new receptor interactions, an important step in cross-species transmission. Key mutations in the spike gene of a CCoV such as CCoV-HuPn-2018, acquired either through recombination or diversifying selection, could facilitate adaptation to a new host’s receptor(s). Here, we provide a further perspective on the recombination history involving the spike gene of CCoV-HuPn-2018, with other members of the *Alphacoronavirus* 1 type II species, and characterize selection pressures across the spike gene, focusing on where such events are in relation to spike functional domains.

## 2. Methods

### 2.1 Sequences and alignments

We collected complete spike gene sequences from *Alphacoronavirus* 1 type II CoVs available in GenBank (accession numbers appear in Supplementary Table S1). Partial spike sequences of *Alphacoronavirus* 1 were excluded from our analysis, as were strains determined from associated publications, to have been be serially passaged (including experimental inoculations of any kind). CCoV type II viruses are currently split into two groups: CCov2a and CCoV2b. Our choice of CCoV2b as our representative CoV from the dog host is based on the fact the most N-terminus subdomain of the spike protein of CCoV2b and TGEV share sequence similarity to CCoV-HuPn-2018, whereas this is not the case for either CCoV2a or FCoV2. Spike domain site positional mapping is based on serotype I feline infectious peritonitis virus (FIPV1; Yang et al. 2020) which is the only *Alphacoronavirus* 1 with a currently available spike protein crystal structure. Although a similar domain map is not available for type II *Alphacoronavirus* 1s, there is sufficient amino acid sequence similarity between type II *Alphacoronavirus* 1s and FIPV to either closely approximate site positions (e.g., for N terminal subdomains) or to more precisely pinpoint site positions (e.g., RBD). Type I and type II *Alphacoronavirus* 1 viruses were not included together in sequence alignments used for selection analyses, because of the divergence between these two types.

We prepared two sets of alignments for comparative sequence analyses. The first of these (set I) included TGEV, CCoV-HuPn-2018, and the CCoV2b strains. This set of sequences was assembled for the analysis of the N-terminus subdomain and we only consider results involving the first 288 aligned amino acid positions (up to and including position 266 of CCoV-HuPn-2018) referred to as the 0-domain in the FIPV structural paper (Yang et al. 2020), and here. The second set (set II) included all strains, and positions downstream of 289, which represents the beginning of the region where FCoV2 and the other sequences share a high degree of sequence similarity. In-frame nucleotide sequences were translated to amino-acids, aligned with MAFFT (Katoh and Standley 2013), and then mapped back to the nucleotide sequences to produce a codon-aware alignment. The resulting alignments were largely gapless, with the exception of two short regions of indels in alignment set I (specifically, the 0-domain). We excluded these regions from positive selection analyses, since uncertain alignment is known to degrade method performance.

### 2.2 Positive selection and recombination

Both alignment sets were screened for recombination with breakpoints identified using GARD (Kosakovsky Pond et al. 2006); set II was also evaluated with RDP5 (Martin et al. 2020) for an additional level of granularity with regard to determining the polarity of sequence exchanges. Each of the resulting GARD fragments served as input to the selection analyses, concomitant with their respective phylogeny, which was inferred using RaxML (Stamatakis 2014) under the GTR+Γ nucleotide substitution model.

We performed site-, branch-, and alignment-level selection tests based on the dN/dS (nonsynonymous / synonymous) ratio estimation as implemented in the HyPhy software package v.2.5.31 (Kosakovsky Pond et al. 2020). We used the MEME method (Murrell et al. 2012) to look for episodic diversifying selection pressure at individual sites across the entire tree (both sets I and II). We tested the CCoV-HuPn-2018 terminal branch for evidence of selection, both overall (some subset of sites along this branch), using the aBSREL (Smith et al. 2015), and BUSTED (Murrell et al. 2015) methods, and at individual sites using the FEL (Kosakovsky Pond et al. 2005) and MEME (Murrell et al. 2012) methods. We modified the FEL and MEME tests to use parametric bootstrap for estimating the null distribution of the likelihood ratio test instead of the asymptotic distribution used in the published tests. This was done to improve the statistical performance of the tests, when the test set comprises a single branch, with the tradeoff of a significant increase in computational cost. We used 100 parametric bootstrap replicates to generate the distribution of the test statistic under the null hypothesis (neutral evolution or negative selection).

### 2.3 Estimating divergence times

#### 2.3.1 Temporal signal

The temporal signal in each GARD partition was assessed using root-to-tip regression in TempEst v1.5.3 (Rambaut et al. 2016) and tip-dating-randomization tests (TDR) (Duchêne et al. 2015). First, ModelFinder (Kalyaanamoorthy et al. 2017) was used in IQTREE-2 (Minh et al. 2020) to identify the best fitting substitution model for each alignment using Bayesian Information Criterion (BIC). Each tree with the best-fitting substitution model was then used as input for root-to-tip regression analysis, where correlation coefficients were calculated using the heuristic residual mean squared function. If a strong temporal signal exists (a linear relationship between genetic distance and sampling time), the correlation coefficient will be positive. For GARD partitions with correlation coefficient greater than 0.1, temporal signal was confirmed using TDR. The R package TipDatingBeast (Rieux et al. 2017) was used to generate ten random permutations of sample dates for each GARD alignment. BEAST2 (Bouckaert et al. 2014) was then used to estimate the evolutionary rate for both alignments with the true sample dates and alignments for each randomized replicate. If the mean clock rate estimate of the alignment with real sample dates fell outside the 95% highest posterior density (HPD) for the randomized date set, temporal signal was deemed sufficient for subsequent analyses.

#### 2.3.2 Model selection

For each alignment that had sufficient evidence of a temporal signal, the fit of combinations of two molecular clock models (strict and uncorrelated relaxed exponential Drummond et al. 2006)) and two demographic models (constant coalescent and Bayesian skyline plot (Drummond et al. 2005)) were assessed using marginal likelihood estimation. For each model tested, marginal likelihood was calculated using PathSampling (Lartillot & Philippe 2006) within the Model-Selection package in BEAST 2 with 12 steps, 1,000,000 MCMC steps with 25% burn-in, and an alpha of 0.3. The average marginal likelihood estimates from two path sampling runs were compared to other model combinations using Bayes Factors (Kass & Raftery 1995).

#### 2.3.3 Discrete trait analysis

The ancestral state of host species (cat, dog, pig, human) was inferred using discrete ancestral trait mapping in BEAST2 (Bouckaert et al. 2014) for each GARD alignment. Bayesian phylogenies were created using 100 million MCMC steps in BEAST2, sampling every 10,000 steps. Trees were summarized using the BEAST2 package TreeAnnotator v2.6.0, discarding 20% of trees as burn-in. Convergence of the MCMC chains was assessed using Tracer v1.7.1(Rambaut et al. 2018) and the effective sample size for each estimated parameter was confirmed to be greater than 200. Phylogenetic trees were annotated using FigTree v1.4.4 (available from http://tree.bio.ed.ac.uk/software/figtree/).

## 3. Results

### 3.1 GARD partition topologies and positive selection

GARD identified two recombinant partitions in the N-terminal subdomains comprising alignment set I, and eight partitions for the remaining regions of the spike gene for alignment set II. The first alignment set I GARD partition included nearly all of the 0-domain (amino acid alignment positions 1-230; position 220 of CCoV-HuPn-2018; FIPV coordinate 235); the second GARD partition from alignment set I overlaps with the downstream onset of sequence similarity between FCoV2, CCoV2b, TGEV and CCoV-HuPn-2018 (which begins at alignment position 289; position 267 of CCoV-HuPn-2018; FIPV coordinate 287; Fig. 1). The phylogeny of GARD partition 1 (Supplementary Fig. S1) includes CCoV-HuPn-2018 as a separate branch intermediately placed between the CCoV2b and TGEV sequences. Phylogenies of the eight GARD partitions for alignment set II (Supplementary Fig. S1) had CCoV-HuPn-2018 in various topological positions including: closely related to one or more CCoV2b sequences (GARD partitions 2, 4, 9), intermediate between dog/cat and pig viruses (GARD partition 3), and as a sister group to FCoV2 sequences (GARD partitions 3, 5, 6, 7, 8). MEME analysis of the 0-domain (alignment set I) did not identify any positively selected sites, whereas the same analysis of alignment set II identified a total of 9 sites at P≤0.05. Two of these sites overlapped with the single branch tests involving CCoV-HuPn-2018 and had moderate Empirical Bayes Factor values (Fig. 1; EBFs: 128, 48 – sites 549 and 619 respectively); these two sites were in the RBD region of the protein (Fig. 1). The remainder of the MEME sites were identified with supportive EBF values on an assortment of CCoV2b and FCoV2 branches. MEME analysis restricted to the CCoV-HuPn-2018 branch, identified a total of 11 positively selected sites, four in the 0-domain (one in the signal peptide, which was identified with SIGNALP-5.0 (Almagro Armenteros et al. 2019)), three in RBD, one in the C-domain, and three in or adjacent to the Connector Domain (CD) (Fig. 1). FEL analysis of the CCoV-HuPn-2018 branch identified four sites, three of which overlapped with the MEME single branch test, and one unique site in RBD (Fig. 1; positive selection statistics summarized in Supplementary Table S2). aBSREL and BUSTED analyses restricted to the CCoV-HuPn-2018 branch did not identify significant evidence of segment-wide selection pressure, however both aBSREL and BUSTED are known to have lower power when the test branch set is small (Murrell et al. 2015; Smith et al. 2015).

**Fig. 1.**
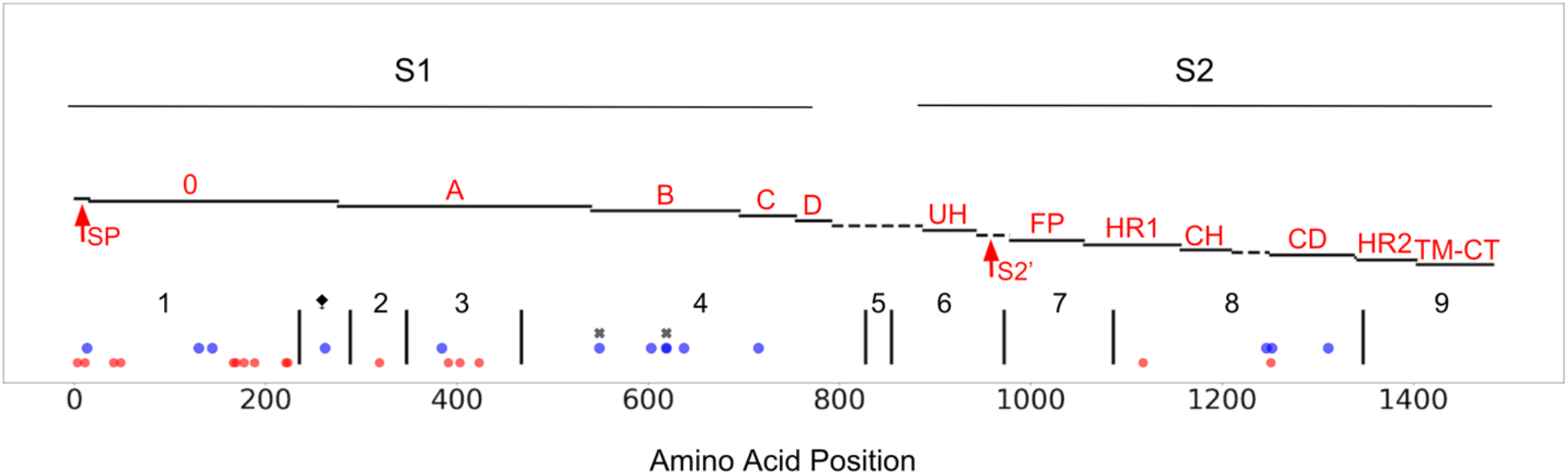
CCoV-HuPn-2018 spike protein, mapped to a published FIPV-UU4 spike gene map (Yang et al. 2020). S1, S2, of spike are highlighted and the protein is further subdivided into functional subunits and subdomains. Blue dots represent sites under positive selection in CCoV-HuPn-2018 as identified by MEME and FEL in the single branch tests; red dots represent sites that are unique in CCoV-HuPn-2018, but are not under positive selection; black “x”s indicate sites under positive selection in the MEME test of the complete alignment that had moderate EBF values for CCoV-HuPn-2018. Red text labels accompany each subdomain/functional unit and are based on the original FIPV spike structure (Yang et al. 2020): SP, signal peptide; 0 domain; A domain; B, includes RBD-Receptor-Binding Domain; C; D; UH, upstream helix; S2’, S2’ cleavage (predicted furin site, using ProP1.0 (Duckert et al. 2004)); FP, fusion peptide; HR1, heptad repeat region 1; CH, central helix; CD, connector domain; HR2, heptad repeat region 2; TM, transmembrane domain; CT, cytoplasmic tail. The dashed line between D and UH refers to a region of peptide with no sequence similarity between FIPV and CCoV-HuPn-2018; this region includes the S1/S2 furin cleavage site in FIPV, which is absent in CCoV-HuPn-2018. The vertical black lines represent the breakpoints of the GARD identified non-recombinant fragments, and are labeled numerically. The ⧪symbol represents a 3’ GARD fragment of alignment set I that was analyzed for positive selection; this GARD fragment was determined from an alignment of just CCoV2b and TGEV sequences (set I). The 5’ end of GARD fragment 2 represents the onset of FCoV2 sequence similarity (set II).

### 3.2 Recombination

RDP5 was used to analyze recombination events in alignment set II which contained FCoV2, CCoV2b, TGEV and CCoV-HuPn-2018 sequences. Because it is not possible to have all natural occurring sequence variants represented in any alignment, it should be noted that when we refer to a sequence as a donor in the following descriptions, it applies to, and is limited by, the sequences in this alignment, and that other closely related genetic variants could be the precise donor. A total of 19 recombination events were well supported (Table S3) by a subset of three or more of the recombination detection methods implemented in RDP5 including BURT, RDP, MaxChi (Smith 1992) and GENECONV (Padidam et al 1999;). Of the 19 supported events, five implicate the CCoV-HuPn-2018 sequence as either a possible recombinant (two events) or as a genetic donor sequence (three events; Supplementary Fig. S2). In the two recombination events where CCoV-HuPn-2018 is the proposed recombinant sequence the putative genetic donors are FCoV2 (strain M91-267, accession AB781788.1) and TGEV (strain TGEV/USA/Tennessee144/2008, accession KX900401.1), and CCoV2b (strain 341/05, accession EU856361.1) and TGEV (strain JS2012, accession KT696544.1), events 13 and 19 respectively. There are three well supported recombination events where CCoV-HuPn-2018 is suggested as the genetic donor, and each event involves a second donor in a strain other than CCoV-HuPn-2018 (i.e., one event with each of FCoV2, CCoV2b, and TGEV respectively). The three inferred recombination events are as follows: CCoV-HuPn-2018 recombines with a FCoV2 (strain Tokyo/cat/130627, accession AB907624.1) sequence to yield a FCoV2 recombinant (strain WSU 79-1683, accession JN634064.1), CCoV-HuPn-2018 recombines with CCoV2b (strain 341/05, accession EU856361.1) to yield a CCoV2b recombinant (strain 174/06, accession EU856362.1), and lastly CCoV-HuPn-2018 recombines with TGEV (strain JS2012, accession KT696544.1) to yield an FCoV2 recombinant (strain WSU 79-1683, JN634064.1), events 5, 6, and 10 respectively. Of the 19 recombination events there are five where the inferred parental sequences and recombinants all belong to different virus types (i.e., TGEV and FCoV2 recombine to yield a CCoV2b) and of these five, one event (event 10) implicates CCoV-HuPN-2018 as a proposed genetic donor, and one (event 13) implicates it as a proposed recombinant.

### 3.3 Temporal analysis

GARD partition 7 had substantive temporal signal in the root-tip-regression and TDR analyses. Four other partitions had correlation coefficients greater than 0.1 on the root-tip-regression tests (Table S4) but failed the TDR, so only GARD partition 7 was used in the temporal analysis (Supplementary Fig. S3). Ancestral host state and lineage divergence times for the GARD 7 partition are shown in Supplementary Fig. S4. CCoV-HuPn-2018 likely diverged from a lineage most recently circulating in cats between 1846 and 1976 (95% HPD – Highest Posterior Density Interval), with a median estimate of 1957. Additional possible host shifts are noted throughout the evolutionary history of the GARD 7 partition including another cat to dog host jump around or after 1981, 95% HPD = [1882, 2005].

## 4. Discussion

We found moderate statistical evidence of positive selection acting upon sites in various regions of the spike gene, primarily at specific sites of the CCoV-HuPn-2018 branch. The 0-domain of CCoV-HuPn-2018 (GARD partition 1; Fig. 1) had seven unique amino acid residues relative to the sequences in the set II alignment, two of which were inferred to be evolving under positive selection (one in the signal peptide). This portion of the CCoV-HuPn-2018 NTD comprises 288 residues with homology only to available TGEV and CCoV2b isolates. Earlier studies have reported on this similarity, implicating recombination between CCoV2 and TGEV (Wesley 1999; Decaro et al. 2009) in the origins of CCoV2b (Wesley 1999). These recombinant CCoV2b are circulating among the domestic dog population (Decaro et al. 2009). The 0-domain region (Fig. 1) is subject to relaxed selection (analysis conducted with RELAX - Wertheim et al. 2015) compared to the other CCoV sequences (GARD partition 1; RELAX results: K=0.07; p=0.005). It has an increased rate of molecular evolution relative to the other CCoV and TGEV sequences (significant Tajima’s relative rate test, implemented in MEGA X – Kumar et al. 2018; all three codon positions, as well synonymous alone, and for most nonsynonymous pairwise comparisons). This selective relaxation suggests that at least some of the history of CCoV-HuPn-2018 involves the loss of function or loosening of functional constraints of this domain, and may also suggest that this region may have different functional roles in humans and dogs.

The role of spike NTD domains in CoV infections is increasingly being recognized. The NTD may act as a co-receptor for SARS-CoV-2, interacting with tyrosine-protein kinase receptor UFO (AXL; Wang et al. 2021) and possibly sialic acids (reviewed in Sun 2021). Sialic acid binding in the NTD has been confirmed for TGEV and PEDV (Schultze et al. 1996; Liu et al. 2015). Point mutations or a short deletion near the N-terminus in TGEV eliminate sialic acid binding, and are associated with lower viral pathogenicity (Krempl et al. 1997). The complete absence of the 0-domain in porcine respiratory coronavirus (PRCV; closely related to TGEV), which includes the region involved in the Krempl et al (1997) experiments, eliminated sialic acid binding (Rasschaert et al. 1990). This 0-domain deletion and the resulting loss in sialic acid binding led to a switch in tropism and pathogenicity for PRCV to predominantly respiratory tract-tropic (Krempl et al. 1997). TGEV on the other hand can infect both the respiratory and enteric tracts (Sanchez et al. 2019). A human respiratory *Alphacoronavirus* - HCoV-229E - hypothesized to have originated in bats, is also missing this region (Corman et al. 2015). Krempl et al. identified, and experimentally confirmed, a ten-residue region of the 0-domain that is essential for TGEV pathogenicity. This same region in our TGEV/CCoV2b/CCoV-HuPn-2018 alignment has three amino acid changes and one amino acid deletion in CCoV-HuPn-2018 relative to TGEV (and no changes in TGEV). Regions upstream and downstream of these ten amino acids are strongly conserved. This general region of the sequence, between 143-168 of our alignment (Supplementary Fig. S5; 143-153 of CCoV-HuPn-2018 sequence), is the most variable section of the 0-domain across these viruses, with two gaps to accommodate two six amino-acid insertions in the CCoV2b sequences. Although these CCoV2b insertions are of the same length, they differ in amino acid composition.

In summary: (a) CCoV-HuPn-2018, shows evidence of relaxed selection in the 0-domain portion of NTD; (b) there are numerous amino acid changes for CCoV-HuPn-2018 in this region that were not associated with signals of positive selection (Fig. 1); (c) the 0-domain of CCoV-HuPn-2018 has an increased rate of molecular evolution; (d) unique amino acid changes, and an amino acid deletion, in CCoV-HuPn-2018 are evident in a ten residue area of the 0-domain experimentally determined to be key to sialic acid binding in TGEV and to affect TGEV pathogenicity (e) this same specific sialic acid binding region of CCoV-HuPn-2018 shares no sequence similarity to CCoV2b, as does much of the surrounding sequence, including two 6 amino acid insertions unique to CCoV2b – a solely enteric pathogen (f) Alpha-respiratory viruses such as PRCV and HCoV-229E have lost the 0-domain; (g) CCoV-HuPn-2018 was associated with a respiratory infection in the Malaysian patients. This leads us to hypothesize that CCoV-HuPn-2018 had an enteric origin, has lost that particular tropism, due in part to mutations in the sialic acid binding region of the 0-domain, resulting in reduced selection pressure for this subdomain. Analogous to other *Alphacoronavirus 1*, such as PRCV, that have lost this region entirely, it has evolved a respiratory tropism. Furthermore, we suggest the possibility that the CCoV-HuPn-2018 lineage might eventually lose this region entirely, just like PRCV, but that we are witnessing early stages of this process. A similar deletion of 197 amino acids has been reported in the 0-domain of some strains of PEDV, an *Alphacornavirus 2* (Oka et al. 2014); in this case, without evidence of tropism shift, but with experimental evidence that this deletion attenuates the virus, likely due to its loss of sialic acid binding (Hou et al. 2017). Vlasova et al. in their analysis of the complete CCoV-HuPn-2018 genome, found that there is a deletion in the middle of the N-protein and a truncated ORF7b compared to other CCoVs and suggested this could have implications in the apparent host shift.

A corollary hypothesis to our ideas on the history of the CCoV-HuPn-2018 0-domain, not mutually exclusive to the one we suggest above, is that the increased genetic variation for this subdomain has somehow pre-adapted the virus to this host shift (see discussion by De Fine Licht 2018), and that the weak signal of positive selection that we observe indicates recent limited host specific adaptation, perhaps reflecting a trajectory of a novel niche expansion. Under such a hypothesis the genetic differences apparent in CCoV-HuPn-2018 expose the virus to new regimes of natural selection, and is analogous to discussions on the role of cryptic genetic variation in facilitating adaptation to new conditions (Paaby and Rockman 2014). Alternatively, the limited positive selection that we do see in this region for CCoV-HuPn-2018 could reflect a remnant of earlier selection in a previous host.

We identified four positively selected sites within the putative RBD of CCoV-HuPn-2018. Based on the FIPV spike structure and the accompanying CoV alignment, only one of these four positively selected sites (at position 619) is in a putative RBD extended loop: specifically, at the end of extended loop 2 (cf. Fig. 4 of Yang et al. 2020). RBD extended loops form the interaction points with the APN receptor in other Alphacoronaviruses (Wong et al. 2017; Yang et al. 2020; Wu et al. 2009) and the specifics of the interaction between these loops and APN, even between closely related viruses, can be very different (Wong et al. 2017).

Another region of note with evidence for positive selection in CCoV-HuPn-2018 was the signal peptide. CoV signal peptides play a role in threading the polypeptide chain through the host endoplasmic reticulum (ER) membrane during protein synthesis. The signal peptide is recognized by the translocon and is pulled through into the ER lumen, with the rest of the polypeptide following. Within the lumen, spike folds and becomes glycosylated. N-linked glycosylation is a common form of viral protein modification, in which a glycan is attached to the amide nitrogen of asparagine at the conserved sequence motif Asn-X-Ser/Thr. Many viruses make use of this host cell process to modify surface proteins, including the spike of CoVs, and this can impact antigenicity and host cell invasion. CoVs have numerous N-linked glycosylation sites in the spike protein, including those structurally verified for the type I *Alphacoronavirus* 1 CoV FIPV (Yang et al. 2020). Recent work on HIV, which like CoV is an enveloped virus, found that the signal peptide can influence the glycan profile and antigenicity of the HIV surface protein gp120 (Yolitz et al. 2018), prompting these authors to suggest that despite the fact the signal peptide is not part of the mature protein, it is likely to be subject to immune pressure. Both the MEME and FEL methods detected positive selection at a codon in the signal peptide of CCoV-HuPn-2018. There are also two additional amino acid changes in the signal peptide of CCoV-HuPn-2018 that differentiate it from other CCoV2b and TGEV sequences. The inferred N-linked glycosylation sites on the CCoV-HuPn-2018 and CCoV2b spike sequences (predicted using NetNglyc;http://www.cbs.dtu.dk/services/NetNGlyc/) while mostly overlapping, have some noteworthy differences: there were 26-29 (mode = 28) N-linked glycosylated sites in the CCoV2b sequences, and there were 29 sites in CCoV-HuPn-2018; of these totals, 3-5 of the CCoV2b sites were in the 0-domain, whereas CCoV-HuPn-2018 had seven in the 0-domain. As a control we also performed the N-linked glycosylation site inference on the FIPV UU4 sequence using NetNglyc and identified almost all of the experimentally verified sites (Yang et al. 2020) for this sequence. Thus, CCoV-HuPn-2018 appears to have a slightly different repertoire of N-linked sites compared to CCoV2b, and with several of these differences occurring in the 0-domain. None of these N-linked glycosylation sites were positively selected in CCoV-HuPn-2018. Positive selection of sites within signal peptides has been reported for other viruses, such as cytomegalovirus, where it was demonstrated that the selected variants affect the timing of signal peptide removal and viral glycoprotein intracellular trafficking (Mozzi et al. 2020). We propose that positive selection in the signal peptide of CCoV-HuPn-2018 could reflect an adaptive role in this new host and that the unique amino acid changes in the signal peptide of CCoV-HuPn-2018 compared to CCoV2b and TGEV, could be playing a role with regards to the N-linked glycan repertoire of this strain.

Given the sample of analyzed sequences there are two recombination events identified by RDP5 where CCoV-HuPn-2018 is the proposed recombinant and three events where CCoV-HuPn-2018, is the proposed major genetic donor of recombinantly transferred sequences (Supplementary Fig. S2). Two of the recombination events identified by RDP5, at the 5’ end of the A-domain (events 5 and 13 in Fig. S2), have similar predicted breakpoint locations which overlap almost exactly with GARD partition 2. For one of these events (event 5), CCoV-HuPn-2018 is identified as a possible sequence donor and FCoV2 as a recombinant, and in the other (event 13), CCoV-HuPn-2018 is the proposed recombinant with FCoV2 as the proposed donor sequence. The detection of both events involved the same FCoV2 sequence (strain WSU 79-1683; accession number JN634064), isolated at Washington State University in 1979 (McKeirnan et al. 1981). A 2009 survey of cats in two Malaysian catteries, using PCR primers designed from strain WSU-79-1683 and FIPV79-1146, found a high prevalence of FCoV test positives (Sharif et al. 2009). While these apparently convergent recombination events could indicate a recombination hotspot, it is also plausible that either (1) there was a single recombination event in FCoV2 that was misidentified as a recombination event in CCoV-HuPn-2018, or (2) there were two unique recombination events, as inferred by RDP5, one in FCoV2 and the other in CCoV-HuPn-2018.

The FCoV2 JN634064 sequence is also a very close sister group to CCoV-HuPn-2018 for GARD partitions 2, 6, 7, and 8 (Supplementary Fig. S1), with 6-8 spanning most of the S2 domain. The BEAST analysis timed the split of WSU 79-1683 and CCoV-HuPn-2018 in partition 7 at 1957. The high prevalence of FCoV2 in Malaysian cats, suggested by the Sharif et al. study, concomitant with their primer construction strategy, suggests the possibility that a WSU 79-1683-like virus could be the prevalent FCoV2 strain in Malaysia. There are also the experimental results of Tresnan et al. (1996) which demonstrate that feline APN can serve as a functional receptor of type II CCoV, TGEV and HCoV-229E, suggesting that cats may act as a mixing vessel for generating recombinant *Alphacoronavirus* 1 CoVs. The origins of WSU 79-1683 may also include two recombination events in the Orf1ab region with FCoV1 and CCoV as sequence donors (Herrewegh et al. 1998). These observations lead us to conclude that WSU 79-1683, or its close relative, has had a prominent role in the evolution of CCoV-HuPn-2018 and that these viruses have repeatedly coinfected hosts, resulting in recombinant progeny.

We propose that at some time in the history of CCoV-HuPn-2018, its spike protein 0-domain may have lost its functional significance. Importantly, other viruses in this group such as PRCV, exhibit a similar evolutionary trajectory where an eventual complete loss of this region of the protein is associated with a shift from enteric to respiratory tropism. Both the molecular details underlying how the loss of this domain contributes to tropic shift of this sort, and the reason(s) that zoonotic host shifts in CoVs are frequently coincidental with respiratory infection, remain important unsolved mysteries. Timing of the origins of CCoV-HuPn-2018 to approximately 1957 suggest that this virus may have been circulating undetected in humans, dogs, cats, or intermediate unidentified hosts for decades. This, in turn, leads us to whole-heartedly concur with the suggestions of Vlasova et al. that a systematic survey should be conducted for the prevalence of CCoV-HuPn-2018 in the host species that comprise the complex history of this virus.

## Conflicts of interest

None declared.

## Acknowledgements

This study received funding (FOA PAR-18-604) from the U.S. Food and Drug Administration’s Veterinary Laboratory Investigation and Response Network (FDA Vet-LIRN) under grant 1U18FD006993-01, awarded to LBG and MJS. SLKP and JDZ were supported in part by grants R01 AI134384 (NIH/NIAID) and U01 GM110749 (NIH/NIGMS). We gratefully acknowledge Jean Millet for helpful advice on *Alphacoronavirus* biology and spike structural domains.

## Notes

### Competing Interest Statement

The authors have declared no competing interest.

