## SupplementalFiles for "Recent zoonotic spillover and tropism shift of a Canine Coronavirus is associated with relaxed selection and putative loss of function in NTD subdomain of spike protein"

### Table S1: Isolate details and sequence accessions for the included CoVs.

| Alignment set | Accession number | Alpha-1 Type | Strain Name | Collection date | Location |
|---|---|---|---|---|---|
| 1 & 2 | MW591993.1 | CCoV-HuPN-2018 | CCoV-HuPN-2018 | 2017 | Malaysia |
| 1 & 2 | EU856361.1 | CCoV2b | 341/05 | Dec 2005 | Italy |
| 1 & 2 | EU924791.1 | CCoV2b | 119-08 | Mar 2008 | Italy |
| 1 & 2 | HQ450377.1 | CCoV2b | 68-09 | 2009 | Greece |
| 1 & 2 | HQ450376.1 | CCoV2b | 66-09 | 2009 | Greece |
| 1 & 2 | EU924790.1 | CCoV2b | 430-07 | Oct 2007 | Italy |
| 1 & 2 | EU856362.1 | CCoV2b | 174-06 | Mar 2006 | Italy |
| 1 & 2 | MT906865.1 | CCoV2b | 2020-7 | 2020 | United Kingdom |
| 1 & 2 | GQ477367.1 | CCoV2b | CCOV-NTU336-2008 | Nov 2008 | Taiwan |
| 1 & 2 | LC190907.1 | CCoV2b | CCOV-Dog-HCM47-2015 | June 2015 | Vietnam |
| 1 & 2 | KX900401.1 | TGEV | TGEV/USA/Tennessee144/2008 | 15 Apr 2008 | USA - Tennessee |
| 1 & 2 | KC609371.1 | TGEV | NA | 21 May 2013 | China |
| 1 & 2 | FJ755618.2 | TGEV | H16 | 1973 | China |
| 1 & 2 | KT696544.1 | TGEV | JS2012 | 2012 | China |
| 1 & 2 | EU074218.2 | TGEV | Attenuated H | NA | China |
| 1 & 2 | X53128.1 | TGEV | TGEV FD772-70 | NA | China |
| 1 & 2 | KX499468.1 | TGEV | TGEV AHHF | 22 Dec 2015 | China |
| 1 & 2 | KC962433.1 | TGEV | TGEV HX | 5 May 2012 | China |
| 1 & 2 | JQ693050.1 | TGEV | NA | NA | South Korea |
| 1 & 2 | KX083668.1 | TGEV | NA | 2015 | China |
| 2 | AB781789.1 | FCOV2 | KUK-H-L | NA | Japan |
| 2 | AB781788.1 | FCOV2 | M91-267 | NA | Japan |
| 2 | AB907624.1 | FCOV2 | Tokyo/cat/130627 | NA | Tokyo, Japan |
| 2 | JN634064.1 | FCOV2 | WSU-79-1683 | 1979 | USA - Washington |

### Figure S1

1.

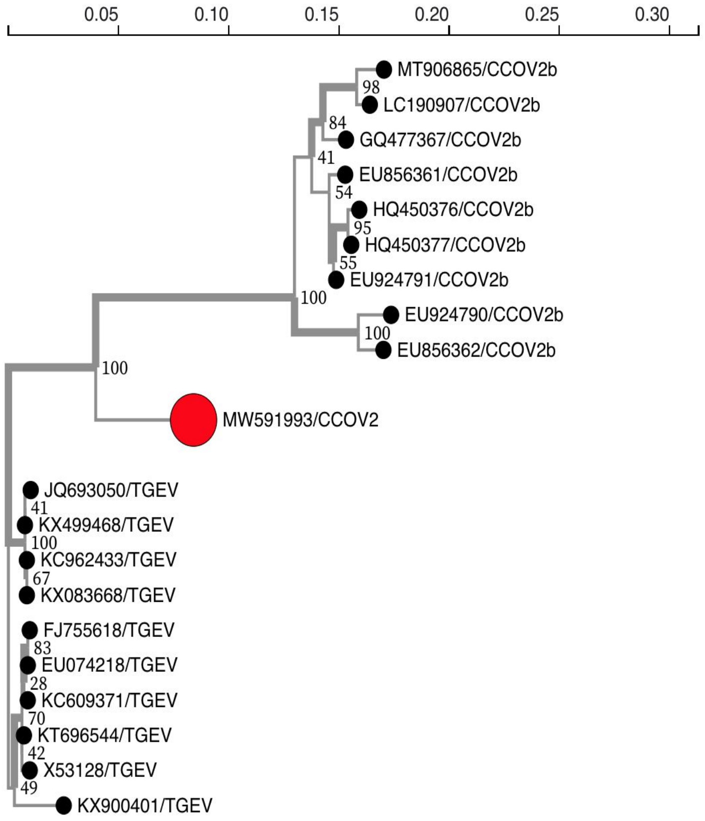
0.05
0.10
0.15
0.20
0.25
0.30
MT906865/CCOV2b
98
LC190907/CCOV2b
84
GQ477367/CCOV2b
41
EU856361/CCOV2b
54
HQ450376/CCOV2b
95
HQ450377/CCOV2b
55
EU924791/CCOV2b
100
EU924790/CCOV2b
100
EU856362/CCOV2b
100
MW591993/CCOV2
JQ693050/TGEV
41
KX499468/TGEV
100
KC962433/TGEV
67
KX083668/TGEV
FJ755618/TGEV
83
EU074218/TGEV
28
KC609371/TGEV
70
KT696544/TGEV
42
X53128/TGEV
49
KX900401/TGEV

2.

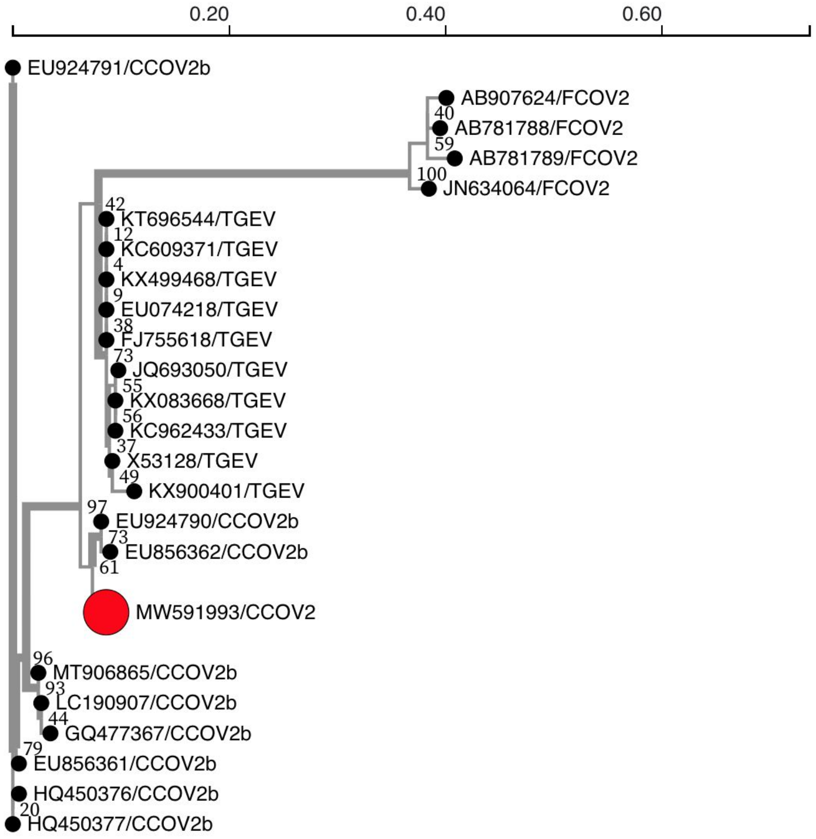
0.20
0.40
0.60
EU924791/CCOV2b
AB907624/FCOV2
40
AB781788/FCOV2
59
AB781789/FCOV2
100
JN634064/FCOV2
42
KT696544/TGEV
12
KC609371/TGEV
4
KX499468/TGEV
9
EU074218/TGEV
38
FJ755618/TGEV
73
JQ693050/TGEV
55
KX083668/TGEV
56
KC962433/TGEV
37
X53128/TGEV
49
KX900401/TGEV
97
EU924790/CCOV2b
73
EU856362/CCOV2b
61
MW591993/CCOV2
96
MT906865/CCOV2b
93
LC190907/CCOV2b
44
GQ477367/CCOV2b
79
EU856361/CCOV2b
HQ450376/CCOV2b
20
HQ450377/CCOV2b

3.

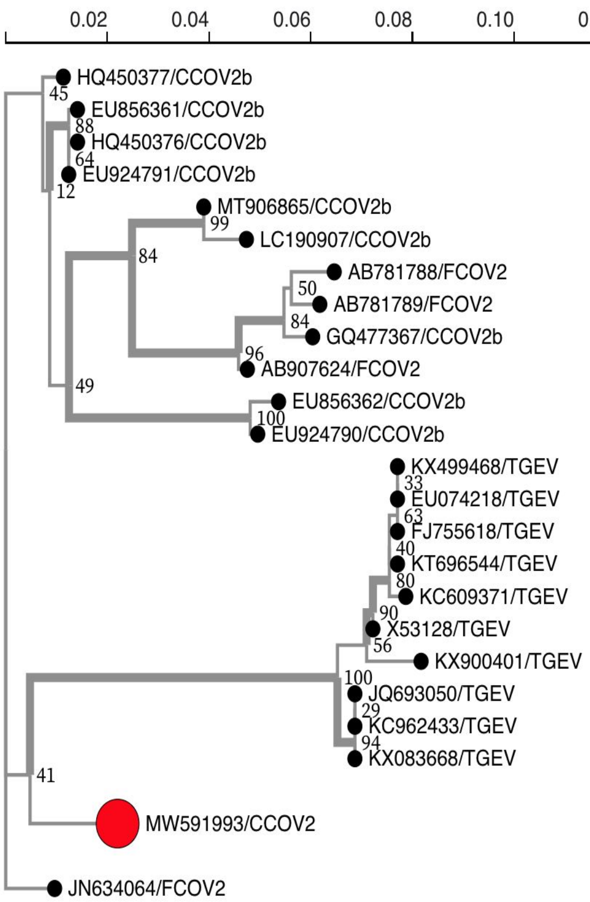

0.02
0.04
0.06
0.08
0.10
0.12
0.14
HQ450377/CCOV2b
45
EU856361/CCOV2b
88
HQ450376/CCOV2b
64
EU924791/CCOV2b
12
MT906865/CCOV2b
99
LC190907/CCOV2b
84
AB781788/FCOV2
50
AB781789/FCOV2
84
GQ477367/CCOV2b
96
AB907624/FCOV2
49
EU856362/CCOV2b
100
EU924790/CCOV2b
KX499468/TGEV
33
EU074218/TGEV
63
FJ755618/TGEV
40
KT696544/TGEV
80
KC609371/TGEV
90
X53128/TGEV
56
KX900401/TGEV
100
JQ693050/TGEV
29
KC962433/TGEV
94
KX083668/TGEV
41
MW591993/CCOV2
JN634064/FCOV2


4.

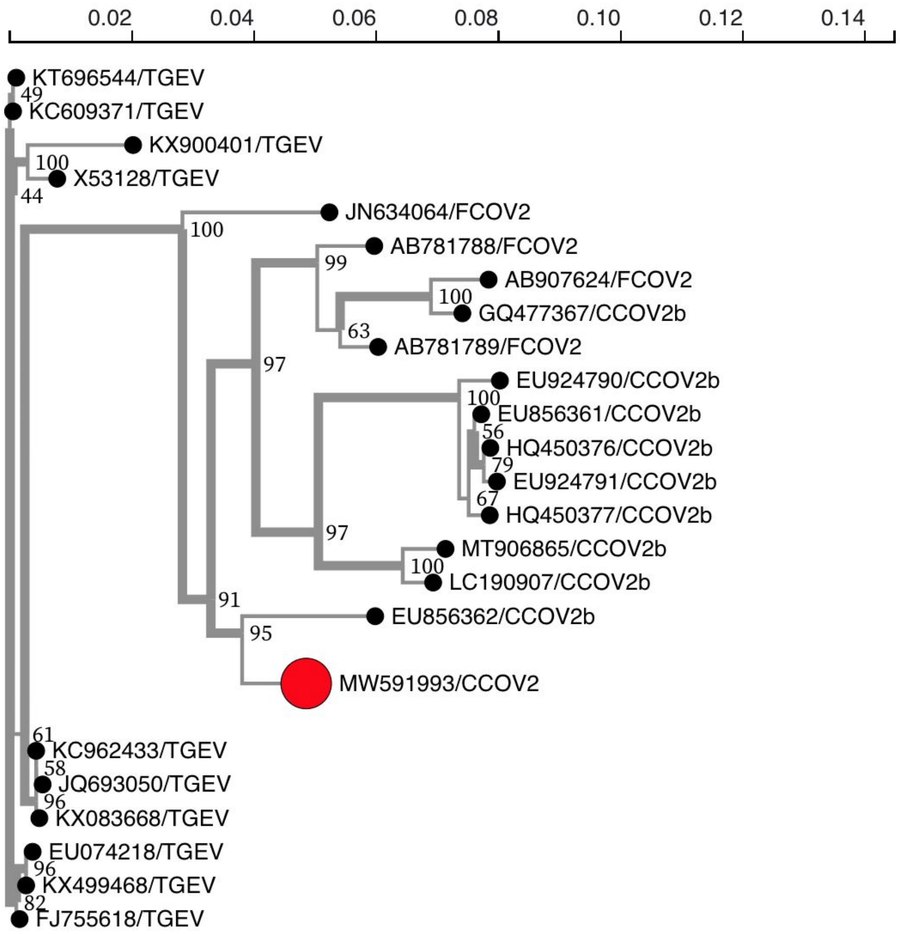

0.02
0.04
0.06
0.08
0.10
0.12
0.14
KT696544/TGEV
49
KC609371/TGEV
KX900401/TGEV
100
X53128/TGEV
44
JN634064/FCOV2
100
AB781788/FCOV2
99
AB907624/FCOV2
100
GQ477367/CCOV2b
63
AB781789/FCOV2
97
EU924790/CCOV2b
100
EU856361/CCOV2b
56
HQ450376/CCOV2b
79
EU924791/CCOV2b
67
HQ450377/CCOV2b
97
MT906865/CCOV2b
100
LC190907/CCOV2b
91
EU856362/CCOV2b
95
MW591993/CCOV2
61
KC962433/TGEV
58
JQ693050/TGEV
96
KX083668/TGEV
EU074218/TGEV
96
KX499468/TGEV
82
FJ755618/TGEV

5.

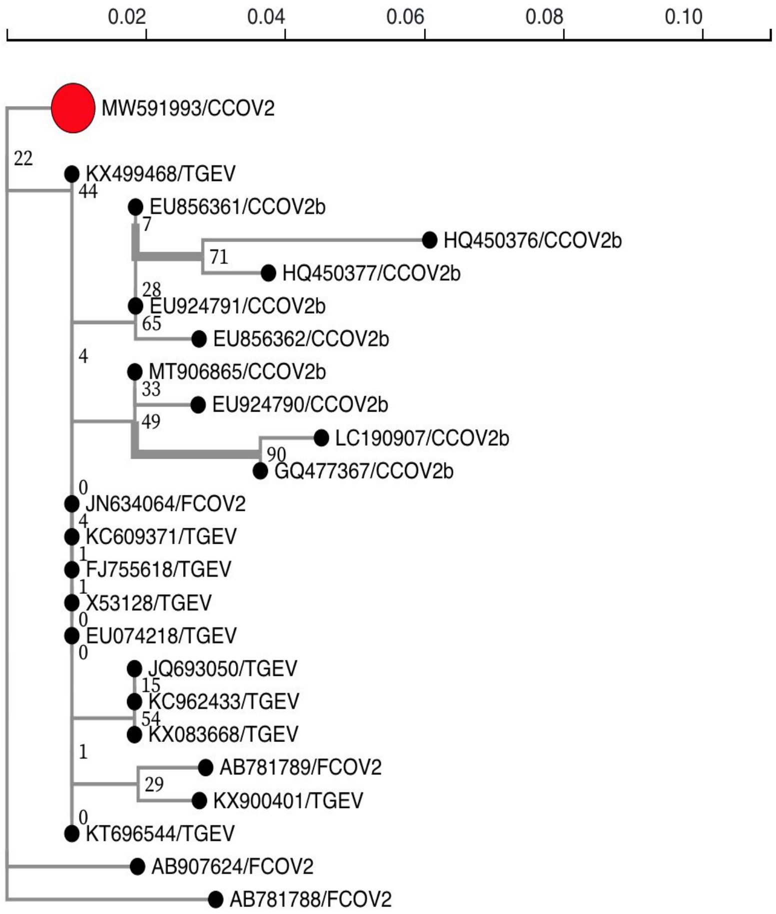

0.02
0.04
0.06
0.08
0.10
MW591993/CCOV2
22
KX499468/TGEV
44
EU856361/CCOV2b
7
HQ450376/CCOV2b
71
HQ450377/CCOV2b
28
EU924791/CCOV2b
65
EU856362/CCOV2b
4
MT906865/CCOV2b
33
EU924790/CCOV2b
49
LC190907/CCOV2b
90
GQ477367/CCOV2b
0
JN634064/FCOV2
4
KC609371/TGEV
1
FJ755618/TGEV
1
X53128/TGEV
0
EU074218/TGEV
0
JQ693050/TGEV
15
KC962433/TGEV
54
KX083668/TGEV
1
AB781789/FCOV2
29
KX900401/TGEV
0
KT696544/TGEV
AB907624/FCOV2
AB781788/FCOV2


6.

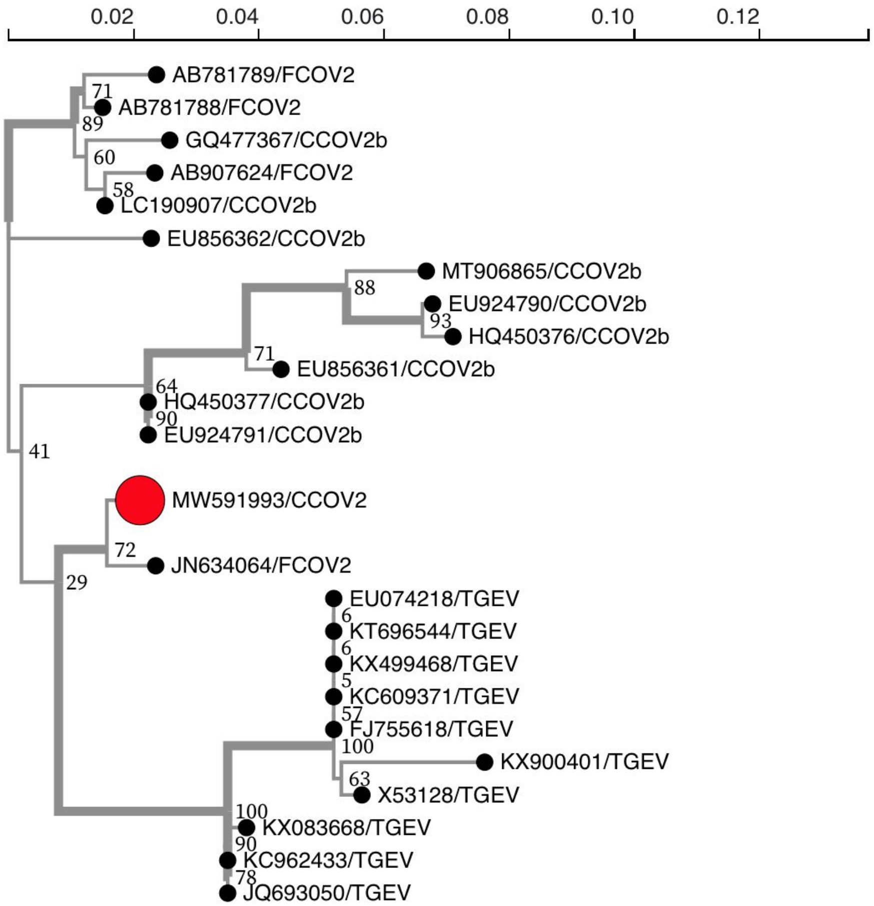

0.02
0.04
0.06
0.08
0.10
0.12
AB781789/FCOV2
71
AB781788/FCOV2
89
GQ477367/CCOV2b
60
AB907624/FCOV2
58
LC190907/CCOV2b
EU856362/CCOV2b
MT906865/CCOV2b
88
EU924790/CCOV2b
93
HQ450376/CCOV2b
71
EU856361/CCOV2b
64
HQ450377/CCOV2b
90
EU924791/CCOV2b
41
MW591993/CCOV2
72
JN634064/FCOV2
29
EU074218/TGEV
6
KT696544/TGEV
6
KX499468/TGEV
5
KC609371/TGEV
57
FJ755618/TGEV
100
KX900401/TGEV
63
X53128/TGEV
100
KX083668/TGEV
90
KC962433/TGEV
78
JQ693050/TGEV

7.

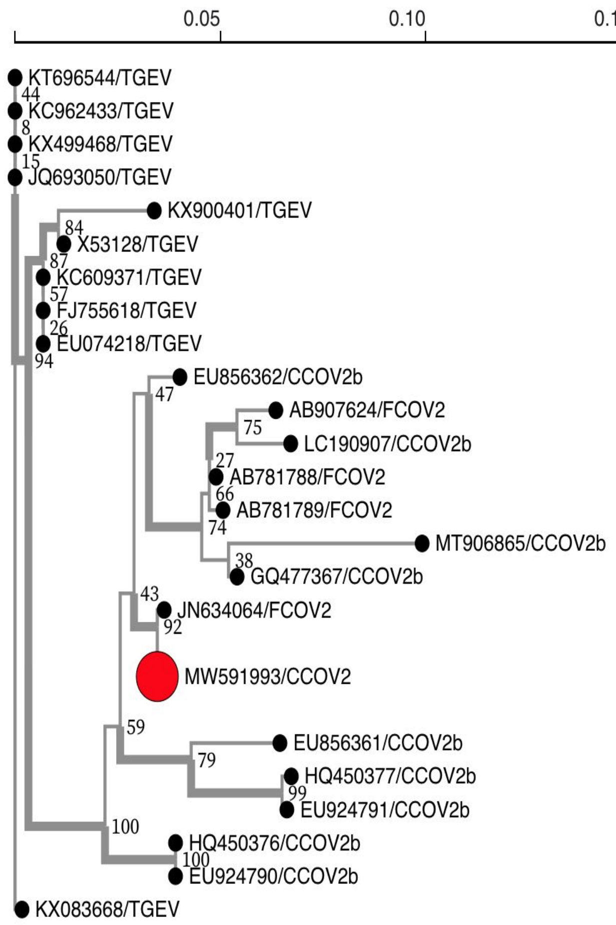
0.05
0.10
0.15
KT696544/TGEV
KC962433/TGEV
KX499468/TGEV
JQ693050/TGEV
KX900401/TGEV
X53128/TGEV
KC609371/TGEV
FJ755618/TGEV
EU074218/TGEV
EU856362/CCOV2b
AB907624/FCOV2
LC190907/CCOV2b
AB781788/FCOV2
AB781789/FCOV2
MT906865/CCOV2b
GQ477367/CCOV2b
JN634064/FCOV2
MW591993/CCOV2
EU856361/CCOV2b
HQ450377/CCOV2b
EU924791/CCOV2b
HQ450376/CCOV2b
EU924790/CCOV2b
KX083668/TGEV

8.

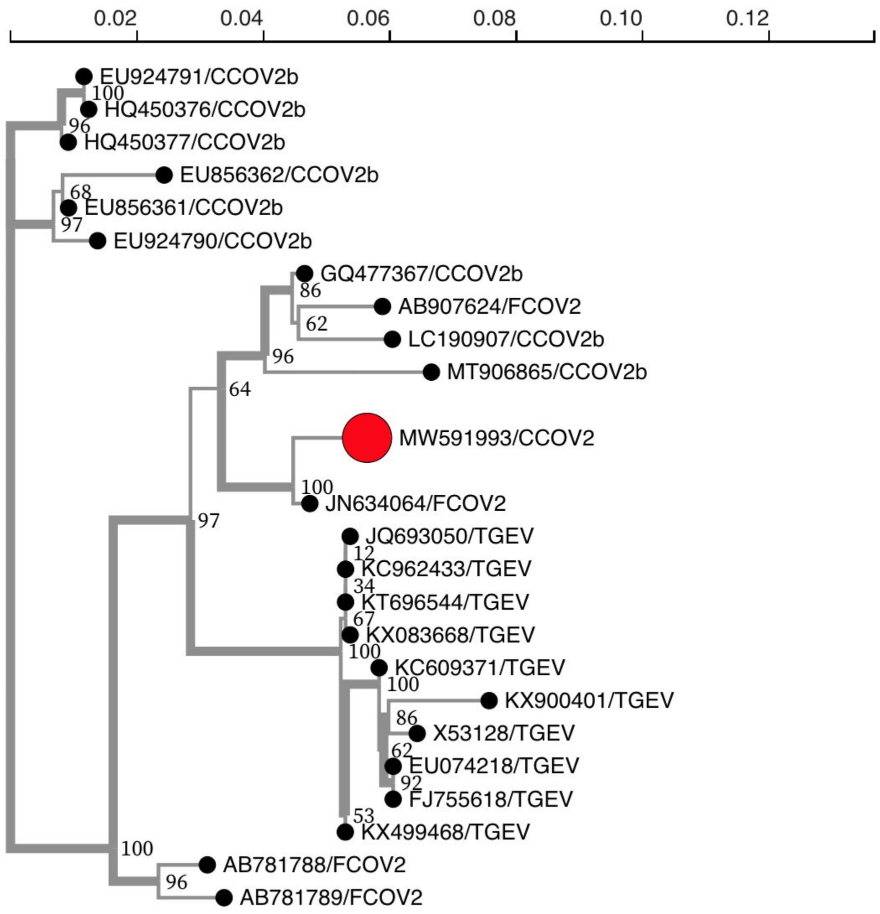
0.02
0.04
0.06
0.08
0.10
0.12
EU924791/CCOV2b
HQ450376/CCOV2b
HQ450377/CCOV2b
EU856362/CCOV2b
EU856361/CCOV2b
EU924790/CCOV2b
GQ477367/CCOV2b
AB907624/FCOV2
LC190907/CCOV2b
MT906865/CCOV2b
MW591993/CCOV2
JN634064/FCOV2
JQ693050/TGEV
KC962433/TGEV
KT696544/TGEV
KX083668/TGEV
KC609371/TGEV
KX900401/TGEV
X53128/TGEV
EU074218/TGEV
FJ755618/TGEV
KX499468/TGEV
AB781788/FCOV2
AB781789/FCOV2

9.

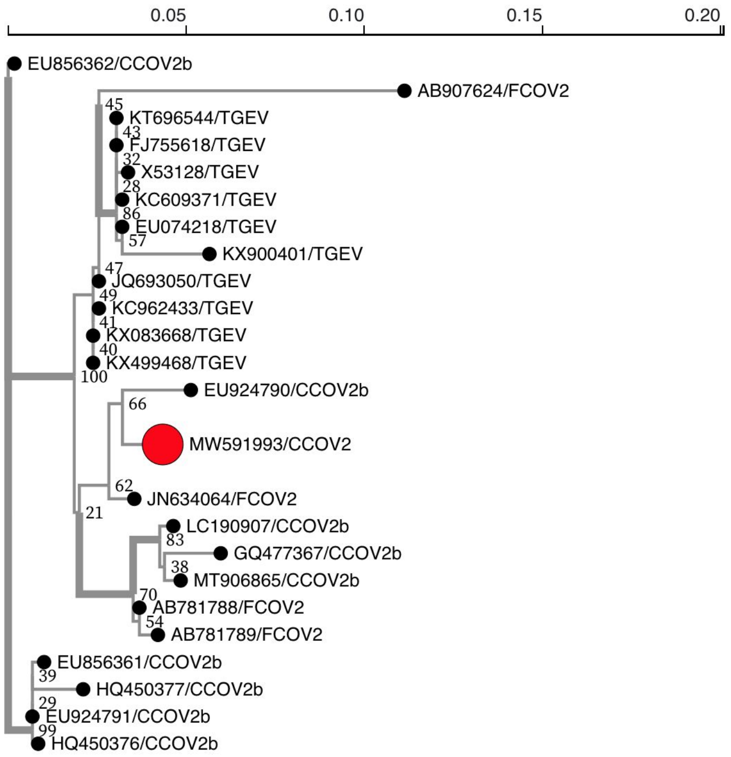


**Figure S1.** Phylogenetic trees for each GARD fragment inferred using RaxML. 1. Is the first non-recombinant fragment from alignment set I and the further trees correspond to non-recombinant fragments from alignment set II. Tree 2 corresponds to non-recombinant fragment 2 as depicted in the Fig. 1 gene map;  tree 3 corresponds to non-recombinant fragment 3, and so on. These trees represent the phylogenetic incongruencies between the different non-recombinant fragments.

### Table S2: Positive selection statistics from MEME and FEL

| HuPN-2018 sites | FIPV mapped sites | MEME Non-synonymous substitution rate | MEME Synonymous substitution rate | FEL Non-synonymous substitution rate | FEL Synonymous substitution rate | MEME p-value | FEL p-value |
|---|---|---|---|---|---|---|---|
| 13 | 13 | 3.58 | 0 | NA | NA | 0.04 | NA |
| 110 | 130 | 30.94 | 0 | 29.432 | 0 | 0.02 | 0.02 |
| 124 | 144 | 3.87 | 0 | NA | NA | 0.02 | NA |
| 246 | 262 | 2.9 | 0 | 2.784 | 0 | 0.02 | 0.04 |
| 367 | 384 | 4.8 | 0 | 4.817 | 0 | 0.04 | 0.05 |
| 575 | 603 | 11.17 | 0 | NA | NA | 0.02 | NA |
| 529 | 549 | 9.23 | 0 | 9.46 | 0 | NA | 0.05 |
| 698 | 715 | 11.26 | 0 | NA | NA | 0.05 | NA |
| 619 | 637 | 12.48 | 0 | 12.47 | 0 | 0.03 | 0.02 |
| 589 | 619 | 10.84 | 0 | NA | NA | 0.04 | NA |
| 1227 | 1246 | 27.49 | 0 | NA | NA | 0.04 | NA |
| 1296 | 1311 | 38.37 | 0 | 37.386 | 0 | 0.01 | 0.01 |
| 1233 | 1252 | 3.3 | 0 | 3.281 | 0 | 0.04 | 0.04 |

### Figure S2

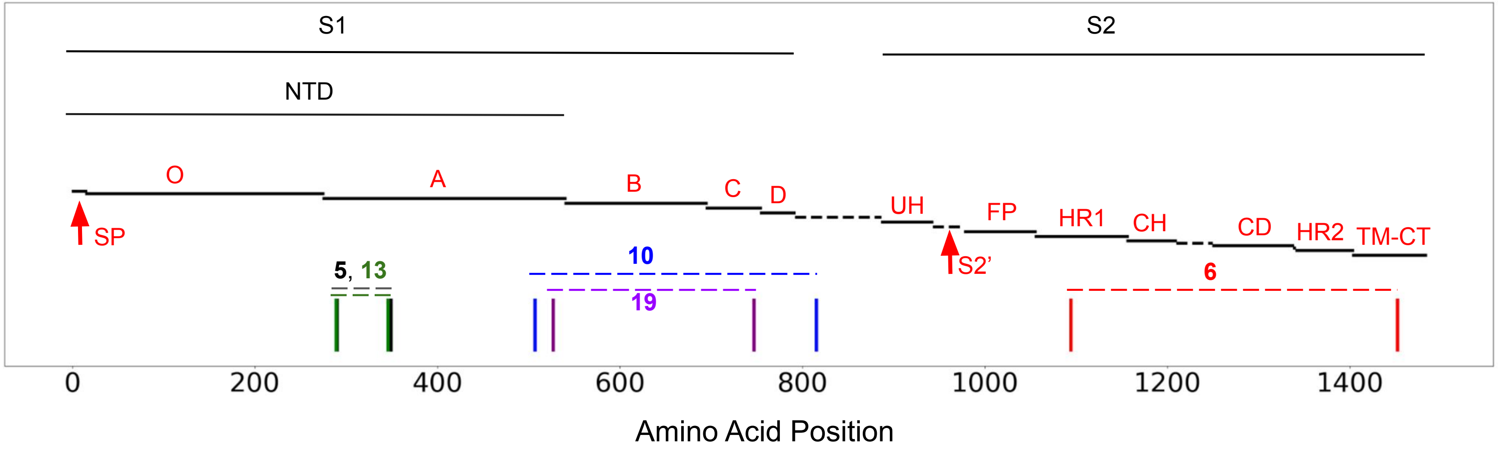


**Figure S2.** RDP5 results with supported recombination events (event boundaries outlined) that implicate CCoV-HuPn-2018, positioned along a spike gene map. A complete, enumerated list of the RDP5 recombination events, involving all sequences, can be found in Table S3. Event 5 (black) involves a proposed FCoV2 recombinant, with genetic donors: CCoV-HuPn-2018 and a different FCoV2. Event 13 (green) extensively overlaps with 5, as well as with GARD fragment 2 (Fig. 1), where CCoV-HuPn-2018 is a proposed recombinant, with genetic donors: FCoV2 from event 13, as well as a TGEV. Event 10 (cyan) proposes the same FCoV2 sequence from 5 and 13 as the recombinant, with genetic donors: CCoV-HuPn-2018 and a TGEV. Event 19 (purple) falls within 10, with CCoV-HuPn-2018 as the proposed recombinant and the genetic donors: a CCoV2b and a TGEV. Event 6 (red) proposes a CCoV2b as the recombinant, with genetic donors: CCoV-HuPn-2018 and another CCoV2b.

### Table S3: RDP5 Recombination Results

**Table key:**

~ = It is possible that this apparent recombination signal could have been caused by an evolutionary process other than recombination.

* = The actual breakpoint position is undetermined (it was most likely either overprinted by a subsequent recombination event or off the edges of the analysed sequence fragments).

^ = The recombinant sequence may have been misidentified (one of the identified parents might be the recombinant)

Minor Parent = Parent contributing the smaller fraction of sequence.

Major Parent = Parent contributing the larger fraction of sequence.

Unknown = Only one parent and a recombinant need be in the alignment for a recombination event to be detectable. The sequence listed as unknown was used to infer the existance of a missing parental sequence.

NS = No significant P-value was recorded for this recombination event using the particular method in question.

| | | Breakpoint Positions | | | | | | |
|---|---|---|---|---|---|---|---|---|
| | | In Alignment | | In Recombinant Sequence | | Relative to AB781789_1_FCOV2_stKUK_H_L_iNA_hFELIS_CATUS_cJAPAN | | |
| Recombination Event Number | Number In .RDP File | Begin | End | Begin | End | Begin | End | Recombinant Sequence(s) |
| 1 | 1 | 1781 | 2398 | 1781 | 2398 | 1781 | 2398 | HQ450376_1_CCOV2b_st66_09_iNA_hDOG_cGREECE_cod2009 |
| 2 | 2 | 3 | 222 | 3 | 222 | 3 | 222 | ^GQ477367_1_CCOV2b_stCCOV_NTU336_F_2008_iNA_hDOG_cT |
| 2 | | | | | | | | |
| 2 | | | | | | | | |
| 2 | | | | | | | | |
| 3 | 3 | 1474 | 3158 | 1474 | 3158 | 1474 | 3158 | LC190907_1_CCOV2b_stCCOV_DOG_HCM47_2015_iNA_hCANIS |
| 4 | 4 | 552 | 1550 | 552 | 1550 | 552 | 1550 | ^EU924790_1_CCOV2b_st430_07_iNA_hDOG_cITALY_codOCT_ |
| 4 | | | | | | | | |
| 4 | | | | | | | | |
| 4 | | | | | | | | |
| 5 | 5 | 3536 | 208 | 3533 | 208 | 3536 | 208 | ^JN634064_1_FCOV2_stWSU_79_1683_iNA_hNA_cNA_codNA_c |
| 6 | 6 | 2426 | 3507 | 2426 | 3507 | 2426 | 3507 | EU856362_1_CCOV2b_st174_06_iNA_hDOG_cITALY_codMAR_ |
| 7 | 7 | 1930* | 3194 | 1930* | 3194 | 1930* | 3194 | ^EU924790_1_CCOV2b_st430_07_iNA_hDOG_cITALY_codOCT_ |
| 8 | 8 | 3538 | 466 | 3538 | 466 | 3538 | 466 | ^MT906865_1_CCOV2b_st2020_7_iNA_hDOMESTIC_DOG_cUNIT |
| 8 | | | | | | | | LC190907_1_CCOV2b_stCCOV_DOG_HCM47_2015_iNA_hCANIS |
| 9 | 9~ | 3380 | 3532* | 3380 | 3532* | 3380 | 3532* | AB907624_1_FCOV2_stTOKYO_CAT_130627_iNA_hFELIS_CAT |
| 9 | | | | | | | | |
| 10 | 10 | 668 | 1546 | 665 | 1543 | 668 | 1546 | ^JN634064_1_FCOV2_stWSU_79_1683_iNA_hNA_cNA_codNA_c |
| 10 | | | | | | | | |
| 10 | | | | | | | | |
| 10 | | | | | | | | |
| 10 | | | | | | | | |
| 10 | | | | | | | | |
| 10 | | | | | | | | |
| 10 | | | | | | | | |
| 10 | | | | | | | | |
| 10 | | | | | | | | |
| 11 | 11~ | 31 | 240 | 31 | 240 | 31 | 240 | EU856362_1_CCOV2b_st174_06_iNA_hDOG_cITALY_codMAR_ |
| 11 | | | | | | | | EU924790_1_CCOV2b_st430_07_iNA_hDOG_cITALY_codOCT_ |
| 11 | | | | | | | | |
| 11 | | | | | | | | |
| 11 | | | | | | | | |
| 11 | | | | | | | | |
| 11 | | | | | | | | |
| 12 | 12 | 21 | 1797 | 21 | 1797 | 21 | 1797 | ^EU856361_1_CCOV2b_st341_05_iNA_hDOG_cITALY_codDEC_ |
| 13 | 13 | 8 | 194 | 8 | 194 | 8 | 194 | ^MW591993_1_CCOV2_stNA_iCCoV_HuPn_2018_hHOMO_cMALAY |
| 14 | 14 | 1810* | 2048 | 1810* | 2048 | 1810* | 2048 | ^MT906865_1_CCOV2b_st2020_7_iNA_hDOMESTIC_DOG_cUNIT |
| 15 | 15 | 2074 | 3542 | 2071 | 3539 | 2074 | 3542 | ^KX499468_1_TGEV_stTGEV_AHHF_iNA_hPORCINE_cCHINA_co |
| 15 | | | | | | | | KT696544_1_TGEV_stJS2012_iNA_hPIG_cCHINA_cod2012_c[T] |
| 16 | 16 | 2488 | 2976* | 2485 | 2973* | 2488 | 2976* | ^AB781788_1_FCOV2_stM91_267_iNA_hFELIS_CATUS_cJAPAN |
| 16 | | | | | | | | AB781789_1_FCOV2_stKUK_H_L_iNA_hFELIS_CATUS_cJAPAN |
| 17 | 17~ | 1* | 212 | 1* | 212 | 1* | 212 | ^HQ450376_1_CCOV2b_st66_09_iNA_hDOG_cGREECE_cod2009 |
| 17 | | | | | | | | EU924791_1_CCOV2b_st119_08_iNA_hDOG_cITALY_codMAR_ |
| 17 | | | | | | | | HQ450377_1_CCOV2b_st68_09_iNA_hDOG_cGREECE_cod2009 |
| 17 | | | | | | | | |
| 17 | | | | | | | | |
| 17 | | | | | | | | |
| 18 | 18 | 208 | 466* | 208 | 466* | 208 | 466* | ^LC190907_1_CCOV2b_stCCOV_DOG_HCM47_2015_iNA_hCANIS |
| 18 | | | | | | | | MT906865_1_CCOV2b_st2020_7_iNA_hDOMESTIC_DOG_cUNIT |
| 18 | | | | | | | | |
| 19 | 19 | 729 | 1393 | 729 | 1393 | 729 | 1393 | MW591993_1_CCOV2_stNA_iCCoV_HuPn_2018_hHOMO_cMALAY |
| 19 | | | | | | | | |
| 19 | | | | | | | | |

~ = It is possible that this apparent recombination signal could have been caused by an evolutionary process other than recombination.

* = The actual breakpoint position is undetermined (it was most likely either overprinted by a subsequent recombination event or off the edges of the analysed sequence fragments).

^ = The recombinant sequence may have been misidentified (one of the identified parents might be the recombinant)

Minor Parent = Parent contributing the smaller fraction of sequence.

Major Parent = Parent contributing the larger fraction of sequence.

Unknown = Only one parent and a recombinant need be in the alignment for a recombination event to be detectable. The sequence listed as unknown was used to infer the existance of a missing parental sequence.

NS = No significant P-value was recorded for this recombination event using the particular method in question.

kpoint Analysis Results

| | | | | | Detection Methods | | | | | |
|---|---|---|---|---|---|---|---|---|---|---|
| Minor Parental Sequence(s) | Major Parental Sequence(s) | RDP | GENECONV | Bootscan | Maxchi | Chimaera | SiSscan | PhylPro | LARD | 3Seq |
| EU924790_1_CCOV2b_st430_07_iNA_hDOG_cITALY_codOCT_ | EU924791_1_CCOV2b_st119_08_iNA_hDOG_cITALY_codMAR_ | 2.54929441781989E-21 | 2.69904252016586E-19 | 1.58842061271349E-21 | 1.60241325509649E-12 | 1.11712519923247E-12 | 3.85798631003641E-13 | NS | NS | 3.09149434769133E-27 |
| HQ450377_1_CCOV2b_st68_09_iNA_hDOG_cGREECE_cod2009 | AB781788_1_FCOV2_stM91_267_iNA_hFELIS_CATUS_cJAPAN | 1.5968974254152E-18 | 1.4763E-15 | NS | NS | 3.42976996758612E-08 | 6.57784854284356E-12 | NS | NS | 1.47659662275146E-13 |
| EU856361_1_CCOV2b_st341_05_iNA_hDOG_cITALY_codDEC_ | | | | | | | | | | |
| EU924791_1_CCOV2b_st119_08_iNA_hDOG_cITALY_codMAR_ | | | | | | | | | | |
| HQ450376_1_CCOV2b_st66_09_iNA_hDOG_cGREECE_cod2009 | | | | | | | | | | |
| AB907624_1_FCOV2_stTOKYO_CAT_130627_iNA_hFELIS_CAT | MT906865_1_CCOV2b_st2020_7_iNA_hDOMESTIC_DOG_cUNIT | 5.71429583963982E-13 | 1.91705875741076E-06 | 1.30171681145963E-11 | 2.17851813857237E-15 | 3.01222470951235E-03 | 3.04126195422575E-08 | NS | NS | 1.22223432622612E-20 |
| HQ450377_1_CCOV2b_st68_09_iNA_hDOG_cGREECE_cod2009 | EU856362_1_CCOV2b_st174_06_iNA_hDOG_cITALY_codMAR_ | 1.06565559083198E-10 | 1.61019545136601E-10 | 2.04958616039796E-10 | 3.81617339524108E-14 | 2.67182105696006E-08 | 8.49132498380889E-15 | NS | NS | 1.47659662275146E-13 |
| EU856361_1_CCOV2b_st341_05_iNA_hDOG_cITALY_codDEC_ | | | | | | | | | | |
| EU924791_1_CCOV2b_st119_08_iNA_hDOG_cITALY_codMAR_ | | | | | | | | | | |
| HQ450376_1_CCOV2b_st66_09_iNA_hDOG_cGREECE_cod2009 | | | | | | | | | | |
| AB907624_1_FCOV2_stTOKYO_CAT_130627_iNA_hFELIS_CAT | MW591993_1_CCOV2_stNA_iCCoV_HuPn_2018_hHOMO_cMALAY | NS | 0.0000000088179 | 1.29024890729025E-07 | NS | 1.61547737751844E-06 | 7.49308129377183E-12 | NS | NS | 2.58640664441145E-09 |
| EU856361_1_CCOV2b_st341_05_iNA_hDOG_cITALY_codDEC_ | MW591993_1_CCOV2_stNA_iCCoV_HuPn_2018_hHOMO_cMALAY | NS | 2.59815477303155E-08 | 8.52400599331166E-13 | 1.34314511108633E-10 | 1.77795291254701E-11 | 2.50494885152511E-15 | NS | NS | 1.47659662275146E-12 |
| EU856361_1_CCOV2b_st341_05_iNA_hDOG_cITALY_codDEC_ | Unknown (EU924791_1_CCOV2b_st119_08_iNA_hDOG_cITALY_codMAR_) | 6.96405652714313E-05 | NS | 1.13700574314452E-04 | 1.82373326519041E-08 | 2.72418943094248E-02 | NS | NS | NS | 2.40326399873259E-04 |
| HQ450377_1_CCOV2b_st68_09_iNA_hDOG_cGREECE_cod2009 | AB781789_1_FCOV2_stKUK_H_L_iNA_hFELIS_CATUS_cJAPAN | 9.06380073189918E-05 | 9.80678186227046E-10 | 2.50077762261688E-12 | 6.53884904347062E-09 | 2.82361660755354E-08 | 1.88070874106544E-18 | NS | NS | 4.36800248615477E-06 |
| Unknown (MT906865_1_CCOV2b_st2020_7_iNA_hDOMESTIC_DOG_cUNIT) | AB781789_1_FCOV2_stKUK_H_L_iNA_hFELIS_CATUS_cJAPAN | NS | 0.000000017955 | 2.93054559450642E-10 | 2.36128641144729E-02 | 5.76594627581528E-03 | 2.37152292533844E-07 | NS | NS | 1.35269529157434E-04 |
| Unknown(LC190907_1_CCOV2b_stCCOV_DOG_HCM47_2015_iNA_hCANIS) | AB781788_1_FCOV2_stM91_267_iNA_hFELIS_CATUS_cJAPAN | | | | | | | | | |
| KT696544_1_TGEV_stJS2012_iNA_hPIG_cCHINA_cod2012_c | MW591993_1_CCOV2_stNA_iCCoV_HuPn_2018_hHOMO_cMALAY | 3.94299401437865E-02 | NS | NS | 1.74085030001778E-05 | 6.39077111921958E-04 | NS | NS | NS | 3.19260417075828E-03 |
| KX900401_1_TGEV_stTGEV_USA_TENNESSEE144_2008_iNA_h | | | | | | | | | | |
| KC609371_1_TGEV_stNA_iZH_hPIG_cCHINA_codNA_crd21_M | | | | | | | | | | |
| FJ755618_2_TGEV_stH16_iNA_hPIG_cCHINA_cod1973_crd1 | | | | | | | | | | |
| EU074218_2_TGEV_stATTENUATED_H_iNA_hNA_cCHINA_codN | | | | | | | | | | |
| X53128_1_TGEV_stTGEV_FS772_70_VIRULENT_BRITISH_ISO | | | | | | | | | | |
| KX499468_1_TGEV_stTGEV_AHHF_iNA_hPORCINE_cCHINA_co | | | | | | | | | | |
| KC962433_1_TGEV_stNA_iTGEV_HX_hSUS_SCROFA_cCHINA_c | | | | | | | | | | |
| JQ693050_1_TGEV_stNA_iDAE_hSWINE_cSOUTH_KOREA_codN | | | | | | | | | | |
| KX083668_1_TGEV_stNA_iHE_1_hSWINE_cCHINA_cod2015_c | | | | | | | | | | |
| KX083668_1_TGEV_stNA_iHE_1_hSWINE_cCHINA_cod2015_c | Unknown (AB907624_1_FCOV2_stTOKYO_CAT_130627_iNA_hFELIS_CAT) | NS | 0.00000015694 | NS | 8.83742902309859E-03 | 3.76911218069598E-04 | NS | NS | NS | NS |
| KX900401_1_TGEV_stTGEV_USA_TENNESSEE144_2008_iNA_h | Unknown(AB781789_1_FCOV2_stKUK_H_L_iNA_hFELIS_CATUS_cJAPAN) | | | | | | | | | |
| KC609371_1_TGEV_stNA_iZH_hPIG_cCHINA_codNA_crd21_M | Unknown(AB781788_1_FCOV2_stM91_267_iNA_hFELIS_CATUS_cJAPAN) | | | | | | | | | |
| KT696544_1_TGEV_stJS2012_iNA_hPIG_cCHINA_cod2012_c | | | | | | | | | | |
| EU074218_2_TGEV_stATTENUATED_H_iNA_hNA_cCHINA_codN | | | | | | | | | | |
| X53128_1_TGEV_stTGEV_FS772_70_VIRULENT_BRITISH_ISO | | | | | | | | | | |
| KX499468_1_TGEV_stTGEV_AHHF_iNA_hPORCINE_cCHINA_co | | | | | | | | | | |
| HQ450377_1_CCOV2b_st68_09_iNA_hDOG_cGREECE_cod2009 | Unknown (EU924791_1_CCOV2b_st119_08_iNA_hDOG_cITALY_codMAR_) | NS | NS | NS | 5.519095479311E-08 | 1.01555418986308E-05 | NS | NS | NS | 1.11866912377856E-06 |
| KX900401_1_TGEV_stTGEV_USA_TENNESSEE144_2008_iNA_h | Unknown (AB781788_1_FCOV2_stM91_267_iNA_hFELIS_CATUS_cJAPAN) | 3.12809262916968E-04 | NS | 5.2554920803031E-06 | NS | NS | 2.26944812454335E-10 | NS | NS | 1.66035638800721E-03 |
| EU856361_1_CCOV2b_st341_05_iNA_hDOG_cITALY_codDEC_ | AB781789_1_FCOV2_stKUK_H_L_iNA_hFELIS_CATUS_cJAPAN | 6.877666015684E-04 | NS | 1.18916131310652E-03 | 5.2541093519971E-03 | 1.63968470554917E-03 | NS | NS | NS | 1.08625952189634E-04 |
| KX083668_1_TGEV_stNA_iHE_1_hSWINE_cCHINA_cod2015_c | EU074218_2_TGEV_stATTENUATED_H_iNA_hNA_cCHINA_codN | NS | NS | NS | 8.23395529029022E-05 | 6.57226784482268E-04 | 3.57411410115915E-04 | NS | NS | 1.86295488697752E-05 |
| | FJ755618_2_TGEV_stH16_iNA_hPIG_cCHINA_cod1973_crd1 | | | | | | | | | |
| EU924791_1_CCOV2b_st119_08_iNA_hDOG_cITALY_codMAR_ | LC190907_1_CCOV2b_stCCOV_DOG_HCM47_2015_iNA_hCANIS | NS | NS | NS | 3.14472616262718E-04 | 3.26398151918505E-03 | 8.30480787757935E-03 | NS | NS | 5.35893306119373E-03 |
| HQ450377_1_CCOV2b_st68_09_iNA_hDOG_cGREECE_cod2009 | AB907624_1_FCOV2_stTOKYO_CAT_130627_iNA_hFELIS_CAT | | | | | | | | | |
| KT696544_1_TGEV_stJS2012_iNA_hPIG_cCHINA_cod2012_c | AB907624_1_FCOV2_stTOKYO_CAT_130627_iNA_hFELIS_CAT | NS | NS | NS | 3.48572332854324E-03 | 1.42536925312427E-03 | NS | NS | NS | NS |
| KX900401_1_TGEV_stTGEV_USA_TENNESSEE144_2008_iNA_h | AB781789_1_FCOV2_stKUK_H_L_iNA_hFELIS_CATUS_cJAPAN | | | | | | | | | |
| KC609371_1_TGEV_stNA_iZH_hPIG_cCHINA_codNA_crd21_M | | | | | | | | | | |
| EU074218_2_TGEV_stATTENUATED_H_iNA_hNA_cCHINA_codN | | | | | | | | | | |
| X53128_1_TGEV_stTGEV_FS772_70_VIRULENT_BRITISH_ISO | | | | | | | | | | |
| KX499468_1_TGEV_stTGEV_AHHF_iNA_hPORCINE_cCHINA_co | | | | | | | | | | |
| AB907624_1_FCOV2_stTOKYO_CAT_130627_iNA_hFELIS_CAT | Unknown (EU074218_2_TGEV_stATTENUATED_H_iNA_hNA_cCHINA_codN) | NS | NS | 4.83472613678042E-02 | 8.68564366541979E-06 | 1.16862405208412E-06 | NS | NS | NS | 2.95064125133138E-07 |
| AB781789_1_FCOV2_stKUK_H_L_iNA_hFELIS_CATUS_cJAPAN | Unknown(FJ755618_2_TGEV_stH16_iNA_hPIG_cCHINA_cod1973_crd1) | | | | | | | | | |
| AB781788_1_FCOV2_stM91_267_iNA_hFELIS_CATUS_cJAPAN | | | | | | | | | | |
| KT696544_1_TGEV_stJS2012_iNA_hPIG_cCHINA_cod2012_c | EU856361_1_CCOV2b_st341_05_iNA_hDOG_cITALY_codDEC_ | NS | NS | NS | 4.76176649908222E-03 | 4.10098289581509E-02 | 6.80442619453942E-19 | NS | NS | NS |
| HQ450377_1_CCOV2b_st68_09_iNA_hDOG_cGREECE_cod2009 | KC609371_1_TGEV_stNA_iZH_hPIG_cCHINA_codNA_crd21_M | | | | | | | | | |
| HQ450376_1_CCOV2b_st66_09_iNA_hDOG_cGREECE_cod2009 | KT696544_1_TGEV_stJS2012_iNA_hPIG_cCHINA_cod2012_c | | | | | | | | | |

### Figure S3

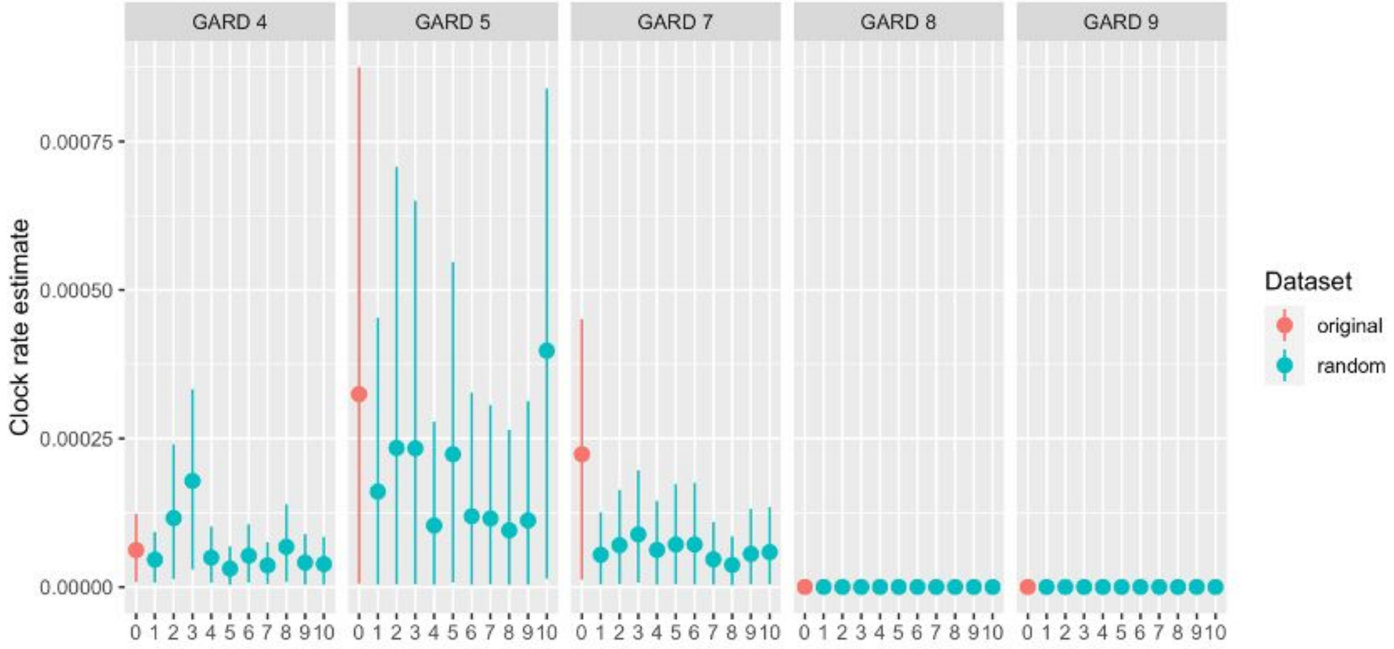


**Figure S3.** Mean and 95% HPD clock rate estimates for the original dataset and ten datasets with random dates are shown for the five GARD fragments with evidence of temporal signal on the root-tip regression analysis. Only GARD 7 had a mean clock rate estimate above the 95% HPD of the randomized datasets, indicating the presence of a temporal signal.

### Table S4: Root-tip-regression results for each GARD Partition

| GARD partition | Correlation coefficient | $R^2$ |
|---|---|---|
| 1 | 0.083 | 0.0070 |
| 2 | -0.046 | 0.0021 |
| 3 | -0.11 | 0.013 |
| 4 | 0.20 | 0.040 |
| 5 | 0.20 | 0.039 |
| 6 | 0.088 | 0.0077 |
| 7 | 0.36 | 0.13 |
| 8 | 0.30 | 0.089 |
| 9 | 0.21 | 0.046 |

### Figure S4

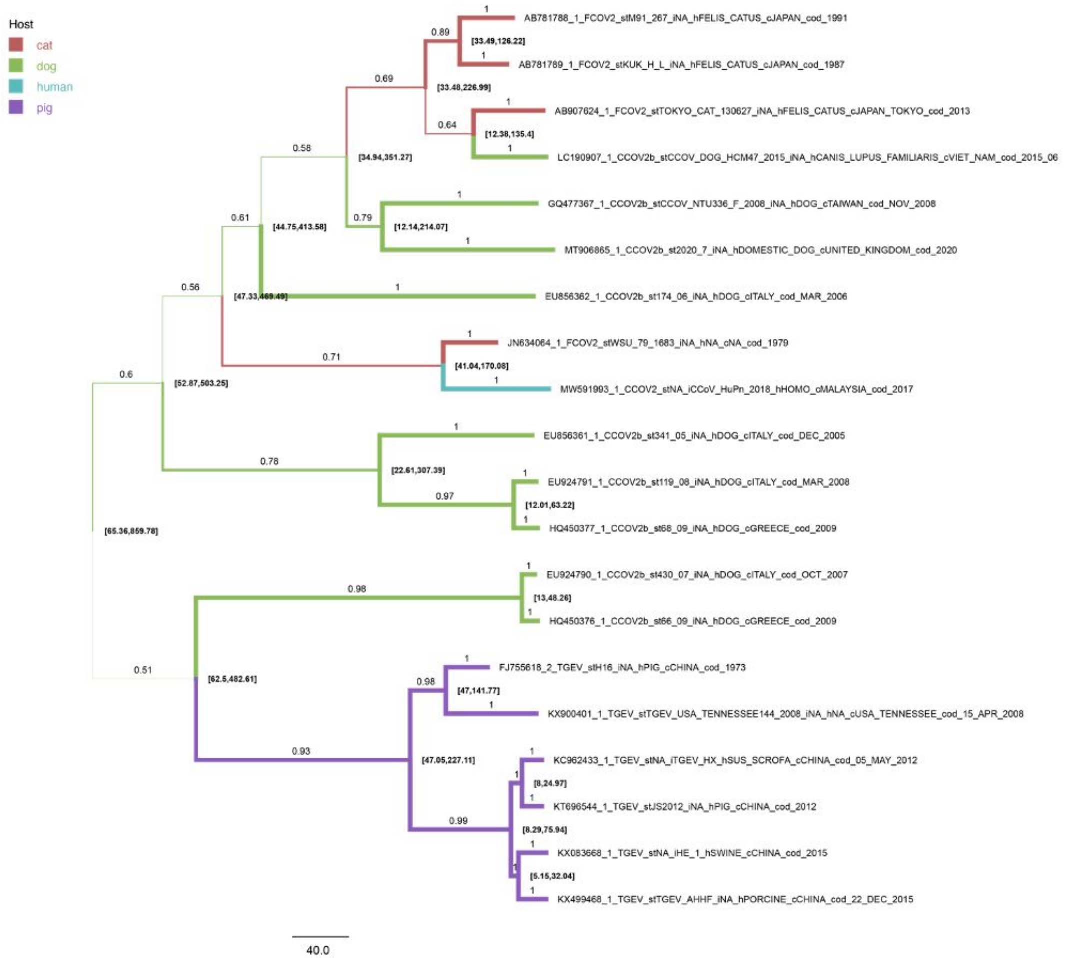
Host
cat
dog
human
pig
AB781788_1_FCOV2_stM91_267_iNA_hFELIS_CATUS_cJAPAN_cod_1991
AB781789_1_FCOV2_stKUK_H_L_iNA_hFELIS_CATUS_cJAPAN_cod_1987
AB907624_1_FCOV2_stTOKYO_CAT_130627_iNA_hFELIS_CATUS_cJAPAN_TOKYO_cod_2013
LC190907_1_CCOV2b_stCCOV_DOG_HCM47_2015_iNA_hCANIS_LUPUS_FAMILIARIS_cVIET_NAM_cod_2015_06
GQ477367_1_CCOV2b_stCCOV_NTU336_F_2008_iNA_hDOG_cTAIWAN_cod_NOV_2008
MT906865_1_CCOV2b_st2020_7_iNA_hDOMESTIC_DOG_cUNITED_KINGDOM_cod_2020
EU856362_1_CCOV2b_st174_06_iNA_hDOG_cITALY_cod_MAR_2006
JN634064_1_FCOV2_stWSU_79_1683_iNA_hNA_cNA_cod_1979
MW591993_1_CCOV2_stNA_iCCoV_HuPn_2018_hHOMO_cMALAYSIA_cod_2017
EU856361_1_CCOV2b_st341_05_iNA_hDOG_cITALY_cod_DEC_2005
EU924791_1_CCOV2b_st119_08_iNA_hDOG_cITALY_cod_MAR_2008
HQ450377_1_CCOV2b_st68_09_iNA_hDOG_cGREECE_cod_2009
EU924790_1_CCOV2b_st430_07_iNA_hDOG_cITALY_cod_OCT_2007
HQ450376_1_CCOV2b_st66_09_iNA_hDOG_cGREECE_cod_2009
FJ755618_2_TGEV_stH16_iNA_hPIG_cCHINA_cod_1973
KX900401_1_TGEV_stTGEV_USA_TENNESSEE144_2008_iNA_hNA_cUSA_TENNESSEE_cod_15_APR_2008
KC962433_1_TGEV_stNA_iTGEV_HX_hSUS_SCROFA_cCHINA_cod_05_MAY_2012
KT696544_1_TGEV_stJS2012_iNA_hPIG_cCHINA_cod_2012
KX083668_1_TGEV_stNA_iHE_1_hSWINE_cCHINA_cod_2015
KX499468_1_TGEV_stTGEV_AHHF_iNA_hPORCINE_cCHINA_cod_22_DEC_2015
[33.49,126.22]
[33.48,226.99]
[12.38,135.4]
[34.94,351.27]
[12.14,214.07]
[44.75,413.58]
[47.33,469.49]
[41.04,170.08]
[52.87,503.25]
[22.61,307.39]
[12.01,63.22]
[65.36,859.78]
[13,48.26]
[62.5,482.61]
[47,141.77]
[47.05,227.11]
[8,24.97]
[8.29,75.94]
[5.15,32.04]
0.89
0.69
0.64
0.58
0.79
0.61
0.56
0.71
0.6
0.78
0.97
0.98
0.51
0.98
0.93
0.99
40.0

**Figure S4.** Ancestral host reconstruction and divergence time estimates**.** Branches are colored by inferred host species. The branch width is proportional to the posterior probability of host assignment (also labeled on branches). Internal nodes are labeled with the divergence time 95% HPD in years from the most recent sample date, 2017.  CCoV-HuPn-2018 diverged from FCoV2 JN634064 between 40 and 170 years ago, with a median date estimate of 1957.

### Figure S5

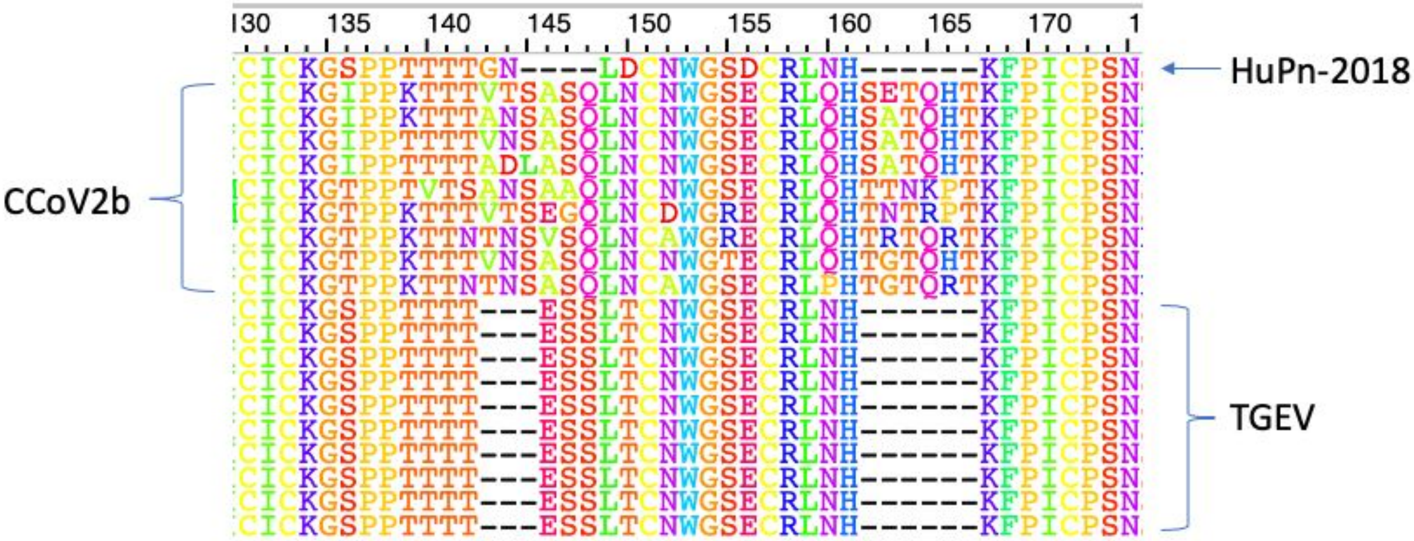


**Figure S5.** Set I sequence alignment highlighting the region involved in the proposed sialic acid binding, identified in the Krempl et al. (1997) mutation experiments.
